# Systemic administration of the mGluR2/3 antagonist LY341495 disrupts reward-related behaviors across ingestive and social domains in mice

**DOI:** 10.64898/2026.04.18.719355

**Authors:** Hiroto Inoue, Mizuki Yamamoto, Yuta Tamai, Shuntaro Matsushima, Kota Yamada, Kazuko Hayashi, Koji Toda

## Abstract

Metabotropic glutamate 2/3 receptors (mGluR2/3) have been implicated in depression, anxiety, learning, and memory, yet their role in reward-related behavior remains poorly understood. Here, we examined the effects of the selective mGluR2/3 antagonist LY341495 on reward-related behaviors in mice using a combination of head-fixed and freely moving behavioral paradigms. In a head-fixed temporal conditioning task, mice developed anticipatory licking and pupil dilation preceding the delivery of a 10% sucrose solution, indicating successful temporal anticipation of reward. Systemic administration of LY341495 dose-dependently reduced both anticipatory and consummatory licking. Normalization analyses further suggested that LY341495 reduced the overall expression of licking behavior while largely preserving its temporal organization. LY341495 also induced pupil dilation and attenuated reward-proximity–related pupillary modulation. To determine whether these effects reflected non-specific motor impairments, we assessed locomotor activity in an open-field task and measured ultrasonic vocalizations (USVs) during courtship interaction. LY341495 did not significantly alter locomotion, excretion, or USV production, suggesting preserved general motor and orofacial motor function. In contrast, LY341495 dose-dependently reduced food intake and decreased social preference, indicating that the effects of mGluR2/3 antagonism extend beyond ingestive behavior to social reward-related processes. These findings demonstrate that mGluR2/3 signaling contributes to the regulation of reward-related behaviors across multiple reward domains independently of general motor dysfunction. These results provide new insight into glutamatergic mechanisms underlying motivation and reward processing and may have implications for neuropsychiatric disorders characterized by anhedonia and motivational deficits.

## Introduction

Glutamate is the principal excitatory neurotransmitter in the mammalian central nervous system and plays a fundamental role in synaptic transmission, neuronal plasticity, and the regulation of complex behaviors. Glutamatergic signaling is mediated by ionotropic glutamate receptors, including α-amino-3-hydroxy-5-methyl-4-isoxazolepropionic acid (AMPA), N-methyl-D-aspartate (NMDA), and kainate receptors, as well as metabotropic glutamate receptors (mGluRs), which are G-protein-coupled receptors that modulate neuronal excitability and neurotransmitter release. Eight mGluR subtypes have been identified and classified into three groups (Group I, Group II, and Group III) based on sequence homology, signal transduction mechanisms, and pharmacological properties (Conn and Pin, 1997; Schoepp and Conn, 1993). Among these, Group II mGluRs, consisting of mGluR2 and mGluR3, are primarily coupled to Gi/o proteins and negatively regulate intracellular signaling through inhibition of adenylyl cyclase activity. Located predominantly at presynaptic terminals, mGluR2/3 function as autoreceptors and heteroreceptors to modulate neurotransmitter release, particularly glutamate transmission (Cartmell and Schoepp, 2000). These receptors are highly expressed in brain regions implicated in emotional regulation and cognitive processes, including the frontal cortex and limbic structures, where they contribute to the regulation of excitatory neurotransmission (Schaffhauser et al., 1998).

mGluR2/3 signaling has been implicated in a broad range of neurological and psychiatric disorders, including depression (Chaki, 2017; Chaki, 2019; Chaki, 2020; Chaki and Watanabe, 2023; Yamamoto et al., 2026), schizophrenia (Li et al., 2015; Dogra and Conn, 2022; Marek, 2010; Maksymetz et al., 2017), addiction (Moussawi and Kalivas, 2010), and anxiety disorders (Schoepp, 2024; Ferraguti, 2018; De Filippis et al., 2015), presumably through their regulatory effects on glutamatergic neurotransmission. Accumulating evidence also suggests that mGluR2/3 contributes to cognitive processes, including learning and memory. However, the effects of mGluR2/3 modulation on cognitive function remain controversial. Higgins et al. (2004) reported that administration of the mGluR2/3 antagonist LY341495 enhanced spatial learning performance in rats in the Morris water maze task. In contrast, Pitsikas et al. (2012) demonstrated that the effects of LY341495 on novel object recognition were dependent on both dose and experimental conditions, suggesting the complexity of mGluR2/3-mediated regulation of cognitive processes. Yoshimizu et al. (2006) showed that repeated administration of the mGluR2/3 antagonist MGS0039 reduced escape failures in the learned helplessness paradigm, suggesting potential antidepressant-like effects of mGluR2/3 blockade. These discrepancies may reflect differences in experimental conditions, including behavioral paradigms, drug doses, and physiological states induced by mGluR2/3 modulation. Therefore, comprehensive evaluation of behavioral and physiological alterations induced by mGluR2/3 modulation may help clarify the mechanisms underlying the diverse effects of mGluR2/3 signaling.

Several studies have suggested that mGluR2/3 blockade enhances dopaminergic signaling in key brain regions involved in reward processing, including the nucleus accumbens and prefrontal cortex. Witkin et al. (2016) demonstrated that administration of LY341495 increased the population activity of spontaneously active dopamine neurons in the ventral tegmental area, elevated extracellular dopamine levels in the nucleus accumbens and prefrontal cortex and potentiated the locomotor-stimulatory effects of the dopamine D2/3 receptor agonist quinpirole. In addition, mGluR2/3 antagonists have been shown to enhance cocaine-induced glutamate release in the nucleus accumbens following chronic cocaine self-administration (Xi et al., 2006) and increase behavioral sensitization in cocaine-treated rats (Yoon et al., 2008). These findings indicate that mGluR2/3 signaling plays an important role in regulating reward-related neural circuits (Moussawi and Kalivas, 2010). However, whether the effects of mGluR2/3 modulation extend beyond drug-related reward processing to influence natural reward behaviors remains largely unknown.

Despite accumulating evidence implicating mGluR2/3 in learning, memory, and reward-related neural processes, the causal contribution of these receptors to natural reward-related behaviors remains poorly understood. A better understanding of the neural mechanisms underlying motivated behaviors requires approaches that allow simultaneous assessment of behavioral performance and physiological states during task execution. In previous studies, we developed a novel head-fixed temporal conditioning task (Toda et al., 2017; Kaneko et al., 2022; Kaneko et al., 2026; Kawai et al., 2022; Ujihara et al., 2024; Yamamoto et al., 2022). This experimental paradigm enables precise measurement of behavioral responses while allowing the recording of physiological parameters, including changes in pupil and eyelid size, during task performance (Kaneko et al., 2022; Yamada and Toda, 2022). This approach provides an opportunity to examine how pharmacological modulation of mGluR2/3 influences not only behavioral outcomes but also associated physiological responses during reward-related behaviors.

In this study, we investigated the effects of LY341495, a selective mGluR2/3 antagonist, on reward-related behavior in mice performing a head-fixed temporal conditioning task. Because licking responses can be influenced by multiple factors, including motor function, motivation, anxiety-related states, and reward processing, we combined this task with complementary behavioral assays to identify the mechanisms underlying LY341495-induced behavioral changes. We first assessed spontaneous locomotor activity and center exploration in a freely moving open-field task to determine whether LY341495 affected general motor function or anxiety-related behavior. We next examined orofacial motor function using a courtship interaction task with ultrasonic vocalization (USV) analysis, as USV production requires coordinated orofacial and respiratory movements. To evaluate motivational processes related to feeding, we measured voluntary food consumption following LY341495 administration. In addition, because reward processing encompasses multiple behavioral domains beyond food-related behaviors, we assessed social preference to determine whether the effects of LY341495 extended to social reward-related behavior. These experiments were designed to distinguish whether LY341495-induced changes in behavioral performance were attributable to nonspecific alterations in motor or physiological states or reflected a broader role of mGluR2/3 signaling in regulating motivational and reward-related processes across ingestive and social domains.

## Methods

### Animals

Male adult C57BL/6J mice were used in the experiment. All mice were naive at the start of the experiment. Mice were maintained on a reversed 12-h light–dark cycle with lights on at 20:00 and temperature controlled (24 ± 2°C), in a humidity of 60 ± 20%. We conducted all the experiments between 13:00 and 20:00. For each subject, we tried to run the experiment at the same period in the day as long as possible. The experimental and housing protocols were approved by the Animal Care and Use Committee of Keio University.

### Surgery

For the mice used for the head-fixed experiments, we conducted surgery to implant a head plate. The mice were anesthetized with 1.0 to 2.5% isoflurane mixed with room air and placed in a stereotactic frame (942WOAE, David Kopf Instruments, Tujunga, CA, USA). The head plate (H.E. Parmer Company, Nashville, TN, USA) was implanted on the skull with a dental cement (Product #56849, 3M Company, Saint Paul, MN, USA) to allow the mice to be head-fixed during the experiment. The mice were group-housed prior to experiments with two to four individuals per cage. After the surgery, the mice were housed singly during recovery for at least two weeks before training began.

### Behavioral Tasks

#### Experiment 1: Temporal conditioning task

To investigate whether the mice can learn to predict the timing of the reward, we trained the mice on a temporal conditioning task with no external sensory cues to signal the timing of the reward (Fig. 1). Eight male mice ranging 4-7 months of age were used in the head-fixed experiment. The mice were water-deprived and received a 10% sucrose solution during the experiments. Their weights were monitored daily. In the head-fixed experiment, we provided the 10% sucrose solution that is enough to maintain over 85% of free-drinking weight of the mice. We also provided additional water provided after the experiment to maintain their weight as needed. The mice had unrestricted access to food in their cages. After recovery from surgery, the mice were water-restricted for two days in their home cage. On the first day of training, the mice were head-fixed briefly and given sucrose rewards to habituate them to the experimental environment. Behavioral experiments were conducted in a square behavioral chamber with a steel drinking spout placed directly in front of the animal’s mouth. Each mouse was kept on a covered elevated platform (custom-designed and 3D printed), with its head fixed by two stabilized clamps holding the sidebars of the head post (Fig. 1A). The heights of the tunnel and clamps were adjusted before each session to ensure comfort and stable data recording. A steel drinking spout was positioned in front of the mouse’s mouth, and both the spout and a meshed copper sheet on the stage were connected to a contact touch-sensor (DCTS-10, Sankei Kizai, Tokyo, Japan) that recorded the timing and duration of licking at 1,000 samples/s. Head-fixed mice were allowed to lick the spout. In the fixed-time schedule task, approximately 2 μL of 10% sucrose solution was delivered through the tube at 10 s intervals (Fig. 1B). Sucrose delivery and recording of licking data were conducted using custom-made Python 3 (version 3.7.7) scripts and a custom-made relay circuit with solenoids. On each day of the experiment, we ran one session that contained 250 trials. One trial consisted of a 10 s interval followed by reward delivery. We defined anticipatory licking as the number of licks from −2 s to 0 s before reward delivery and consummatory licking as the number of licks from 0 s to +2 s after reward delivery.

**Fig. 1:**
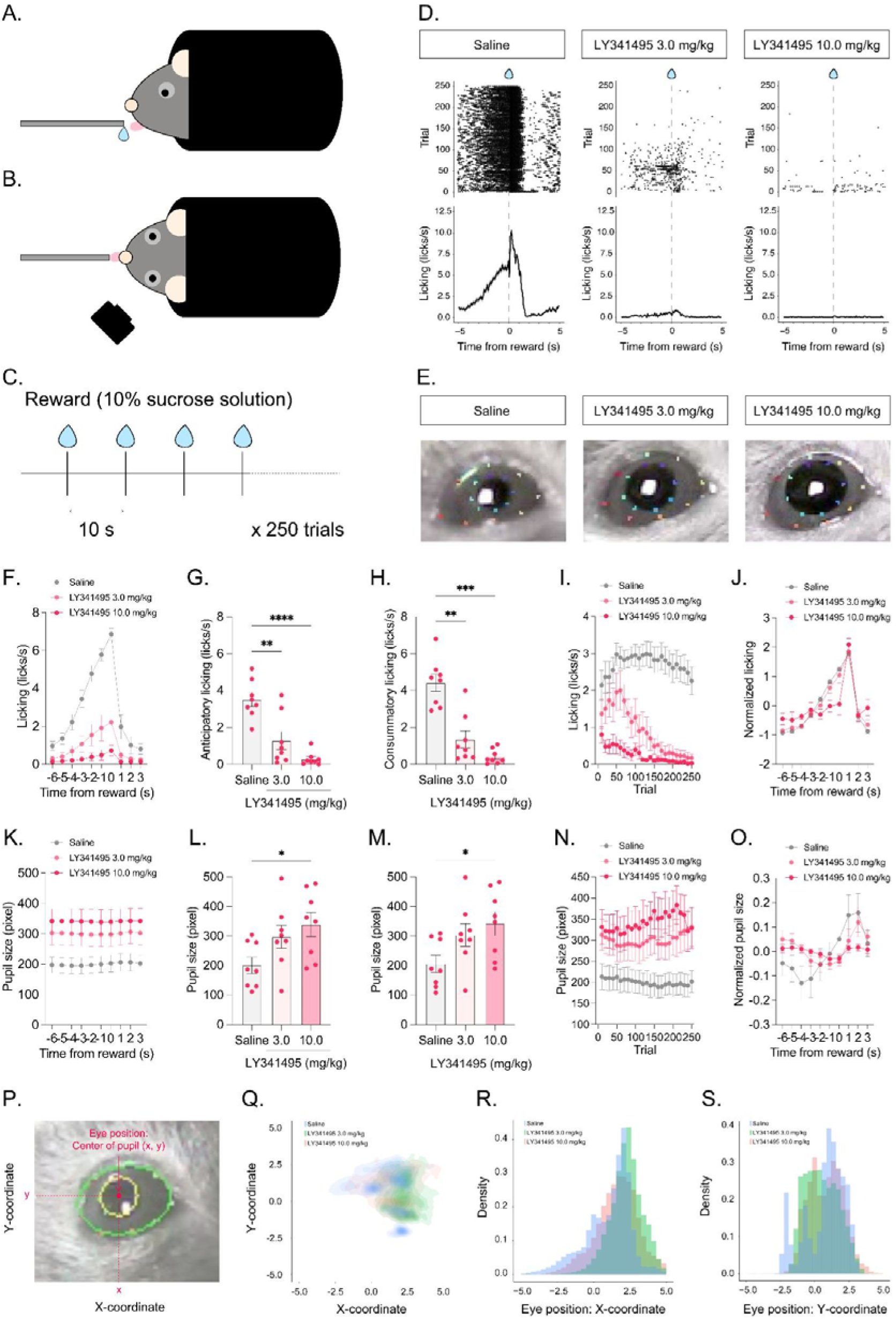
Head-fixed temporal conditioning task. **A.** A schematic illustration of the head-fixed mice viewed from the side. **B.** A schematic illustration of the head-fixed mice viewed from the top. **C.** Temporal conditioning task. In this temporal conditioning task, we delivered a 10% sucrose solution every 10 s with a blunt-tipped needle placed within the licking distance of head-fixed mice. D. Example of the effects of intraperitoneal injections of LY341495 on licking responses. Upper panel shows the raster display of the licking response. Lower panel shows the density plot. Example data of one mouse after the injections of saline, LY341495 3.0 mg/kg, 10.0 mg/kg are indicated in left, center, and right panels, respectively. E. Example data of the effects of intraperitoneal injections of LY341495 on pupil sizes. Saline, LY341495 3.0 mg/kg, 10.0 mg/kg are indicated in left, center, and right panels, respectively. F. Averaged time course of licking responses. G. Averaged time course of pupil dynamics. H. Anticipatory licking. I. Pupil size during the anticipatory period. J. Consummatory licking. K. Pupil size during the consummatory period. L. Licking responses along with trial progress. M. Pupil dynamics along with trial progress. N. Normalized licking response. O. Normalized pupil dynamics. P. Quantifications of eye position. Q. Eye position. R. Distributions of eye position: x-coordinate. S. Distributions of eye position: y-coordinate. \**p* < 0.05, \*\**p* < 0.01, \*\*\**p* < 0.001, \*\*\*\**p* < 0.0001. Error bars represent standard error of the mean.

An infrared camera was used to capture the eyes of mice while the mice performed the fixed-time schedule task. The camera was positioned at 40° from the midline of the mouse and 45 mm from the top of the head. Infrared light was used to capture the pupil. The brightness inside the experimental box was adjusted to 95 lux. DeepLabCut, a tracking tool that uses deep learning, was used to analyze the pupil size (Mathis et al., 2018). First, we trained DeepLabCut to learn the locations of the upper, lower, left, and right edges of the eyelids and edges of the pupil, and the trained model was used to analyze the videos and quantify eyelid and pupil sizes. The median length between the upper and lower eyelids was divided by the median length between the left and right eyelids, and the value multiplied by 10 was quantified as the normalized eyelid size in every frame.

#### Experiment 2: Open-field task

To examine the effects of LY341495 on spontaneous locomotor activity, we conducted an open-field task using nine adult male C57BL/6J mice with the age of 2-4 months. The procedure for the open-field task was adapted from our previous studies (Kaneko et al., 2022; Kaneko et al., 2026; Tamura et al., 2024; Ujihara et al., 2024; Yamamoto et al., 2026). The apparatus was a custom-made white vinyl chloride box measuring 50 cm in length × 50 cm in width × 50 cm in height. Cameras (Logicool HD Webcam C920r, Logicool Co Ltd., Tokyo, Japan) were placed 106 cm above the bottom of the box. The video was recorded on a Windows PC at 30 frames/s. Habituation to the apparatus and saline injection were performed for 60 minutes per day for three days prior to the start of the experiment. During the habituation phase, the animals were inserted into the open field 60 minutes after receiving the intraperitoneal injection of saline and allowed to explore freely for 90 minutes. During the test phase, the order of the saline and LY341495 1, 3, and 10 mg/kg injections was randomized and counterbalanced across subjects. We set more than two days of the intervals between each session of the open-field experiments to avoid the effect of LY341495 injection continuing until the next session. White noise (75 dB) was presented throughout the experiment to mask the external noise. Every time after running the open-field experiment, we wiped inside the box with 70% alcohol and waited 30 minutes for it to dry. We used an open-source visual programming framework, Bonsai (Lopes et al., 2015) to track the locomotor activity of mice using computer vision analysis. The videos were converted to grayscale, smoothened, and monochrome inverted. The mice were identified by setting a contrast threshold. The locomotor activity of the mice in the box was quantified by measuring the changes in the central coordinates of the mice.

#### Experiment 3: Courtship interaction task with recording USVs

We used a male-female courtship interaction task to examine the effects of LY341495 on orofacial movement indexed by USVs of male mice toward female mice. The procedures for the courtship interaction task and USV recording were adapted from our previous studies (Tamura et al., 2024; Yamamoto et al., 2026). We used 12 adult male C57BL/6J mice with the age of 3-6 months. For habituation, adult male mice were paired with female mice in standard cages for at least 2 days to provide sexual experience prior to testing. Males were then habituated to the experimental cage (width 28.0 cm × depth 9.5 cm × height 30.0 cm) for at least 1 day. Subjects received an injection of saline, LY341495 3 and 10 mg/kg 1hour before the test.

During the experiment 3 test a novel female mouse was introduced into the basket, and USVs and approach behavior were recorded for 10 min. A condenser microphone (CM16/CMPA, Avisoft Bioacoustics, Glienicke,Germany) was positioned 31 cm above the floor of the experimental box to record USVs. The recorded sound data was stored on a computer through an audio interface (UltraSoundGate 116Hb, Avisoft Bioacoustics, Glienicke,Germany) using a sampling rate of 96 kHz at 16 bits/sample. A camera (C980GR, Logicool Co Ltd., Tokyo, Japan) was placed next to the microphone and used to track the location of the subject male mouse. All experiments were conducted inside a sound-attenuating chamber (ENV-018V, Med Associates Inc., St. Albans, VT, USA) to minimize external noise.

#### Experiment 4: Food consumption task

To examine the effects of LY341495 on feeding behavior, we conducted a food consumption task using 12 adult male C57BL/6J mice with the age of 2-4 months. The apparatus was a custom-made white vinyl chloride box measuring 50 cm in length × 25 cm in width × 25 cm in height. Cameras (Logicool HD Webcam C920r, Logicool Co Ltd., Tokyo, Japan) were placed 93 cm above the bottom of the box. The video was recorded on a Windows PC at 30 frames/s. Mice were restricted to access to foods in their home cage for 24 hours until the start of the experiment. For each session, mice received an intraperitoneal injection of saline or drug and were returned to their home cages. Sixty minutes after the injection, animals were placed in the test box and allowed to freely explore the apparatus for 10 min for habituation. Following habituation, mice were briefly removed from the box while a piece of waffle was placed inside. The animals were then returned to the apparatus, and food consumption was assessed during a 20-min test period. The order of the saline, LY341495 3 and 10 mg/kg injections was randomized and counterbalanced across subjects. We set more than two days of the intervals between each session of the food consumption experiments to avoid the effect of LY341495 injection continuing until the next session. White noise (75 dB) was presented throughout the experiment to mask the external noise. Every time after running the food consumption experiment, we wiped inside the box with 70% alcohol and waited 30 minutes for it to dry. We used an open-source visual programming framework, Bonsai (Lopes et al., 2015) to track the locomotor activity of mice using computer vision analysis. The videos were converted to grayscale, smoothened, and monochrome inverted. The mice were identified by setting a contrast threshold. The locomotor activity of the mice in the box was quantified by measuring the changes in the central coordinates of the mice.

#### *Experiment 5:* Social preference task

To examine whether mGluR2/3 antagonists affect social reward seeking behavior, we conducted a social preference task. The procedure for the social preference task was adapted from our previous studies (Nasukawa et al., 2022). We used 9 adult C57BL6J male mice with the age 2-3 months. A custom-made white vinyl chloride box (25 cm length × 50 cm width × 30 cm height) was used for the test. Male mice were placed in a white cylindrical pen stand (8 cm diameter × 10 cm height). The pen stands containing the stimulus individuals were placed at the center of both ends of a 50 cm wide box. A 500 mL empty glass bottle was placed on top of the pen stand to prevent the subject mouse from climbing. Two partition walls (5 cm length × 5 mm thickness × 30 cm height) were attached to both sides of the center of the 50 cm width box to separate the spaces on the left and right sides. The cameras (Logicool HD Pro Webcam C920n, Logicool Co Ltd., Tokyo, Japan) were placed 85 cm above the bottom of the box. During the social preference test, the mice were video-recorded on a Windows PC. Mice were intraperitoneally injected with 3 or 10 mg/kg of LY341495 or saline and then placed in the box 60 min after the injection. After 15 min of habituation, the mice were temporarily removed from the box, and the stimulus individuals, male C57BL/6J mouse and a novel object, were placed in the basket. The subject mice were then placed again in the center of the box and allowed to explore freely in the box for 10 min. White noise (75 dB) was presented throughout the experiment to mask the external noise.

### Drug

We obtained LY341495 from Santa Cruz Biotechnology, Inc (Dallas, TX, USA). LY341495 was dissolved in a saline solution. We administered LY341495 to mice via intraperitoneal injection at the dose of 0.01 mL/g. The doses used in the present study were selected based on previous studies reporting behavioral effects of LY341495 in rodents (Higgins et al., 2004; Pałucha-Poniewiera et al., 2021; Pitsikas et al., 2012; Podkowa et al., 2016; Witkin et al., 2016; Zanos et al., 2019) and on preliminary observations from the open-field experiment. Because the 1 mg/kg dose produced minimal behavioral effects in the open-field task, subsequent experiments focused on the 3 and 10 mg/kg doses to evaluate the effects of mGluR2/3 antagonism across multiple behavioral paradigms. The concentrations of 1-10 mg/kg of LY341495 are within a range of commonly used dosage (0.3-10 mg/kg). All the experiments were initiated 60 minutes after the injection of saline or LY341495.

### Analysis

All data analyses were performed using RStudio (version 2022.02.0, RStudio PBC, MA, USA), MATLAB (R2020a, The MathWorks, Inc., Natick, MA, USA) and GraphPad Prism (version 10.3.1, GraphPad, CA, USA). We used DeepLabCut (Mattis et al., 2018) to analyze the pupil size in the head-fixed experiments. In freely moving experiments, we used bonsai (Lopes et al., 2015) to quantify locomotor activity. Video recordings were converted to grayscale, smoothed, and inverted to enhance contrast, and mice were detected by applying a contrast threshold. Locomotor activity was quantified by calculating changes in the animals’ central coordinates. In the head-fixed temporal conditioning experiment, we normalized the pupil diameter by using a Scale() function. It calculates as follows.

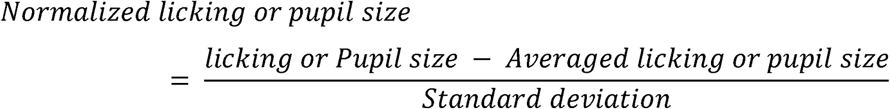

In the USV experiment, we defined mice as spending time with the social interaction if they were located within 3.5 cm of the edge of the box that contained the stimulus mice. To investigate the characteristics of USVs, VocalMat (Fonseca et al., 2021), a machine learning-based platform that enables segmentation of the USVs from the background noise and classifies USVs into distinct call categories, was used.

Sample sizes were not determined by an a priori power analysis. Instead, group sizes were chosen based on our previous studies using similar behavioral paradigms and on sample sizes commonly used in the field. All animals that completed the experiments and met the predefined inclusion criteria were included in the analyses.

## Results

### Experiment 1: temporal conditioning task in head-fixed mice

To investigate whether the mice can learn to predict the timing of the reward, we trained the mice on a temporal conditioning task with no external sensory cues to signal the timing of the reward. In this temporal conditioning task, we delivered a 10% sucrose solution every 10 s with a blunt-tipped needle placed within the licking distance of head-fixed mice (Fig. 1A-C). After the training, all mice showed ramping anticipatory licking toward the timing of reward delivery (Fig. 1D). These results replicated our previous findings (Kaneko et al., 2022; Kaneko et al., 2026; Kawai et al., 2022; Toda et al., 2017; Ujihara et al., 2024; Yamamoto et al., 2022).

To examine the role of mGluR2/3 on the performance of the temporal conditioning task, we examined the effects of intraperitoneal injections of the mGluR2/3 antagonist LY341495 on licking, pupil, and eye position (Fig. 1D,E). LY341495 decreased licking in a dose-dependent manner (Fig. 1F). Both anticipatory and consummatory licking were dose-dependently decreased (anticipatory licking: Fig. 1H, *F* (1.523, 10.66) = 27.34, *p* = 0.0001, repeated-measures one-way ANOVA; Saline v.s. LY341495 3.0 mg/kg: *p* = 0.0069; Saline v.s. LY341495 10.0 mg/kg: *p* < 0.0001 post-hoc Tukey test; consummatory licking: Fig. 1J, *F* (1.916, 13.41) = 40.03, *p* < 0.0001, repeated-measures one-way ANOVA; Saline v.s. LY341495 3.0 mg/kg: *p* =0.0014; Saline v.s. LY341495 10.0 mg/kg: *p* = 0.0001; post-hoc Tukey test). Effects of LY341495 on licking response increased along with trial progress (Fig. 1I). Normalized licking response shows that the reward prediction was not affected in LY341495 3.0 mg/kg condition but disrupted in LY341495 10.0 mg/kg condition (Fig. 1J).

While the mice performed the temporal conditioning task, we recorded a movie of the mice in the head-fixed condition. Using DeepLabCut (Mathis et al., 2018) and eclipse fitting (Yamada & Toda, 2022), we successfully quantified the size of their pupils (Fig. 1E). The pupil sizes of the mice during the temporal conditioning task increased in a dose-dependent manner (Fig. 1K). Pupil size in both anticipatory and consummatory periods were dose-dependently increased (anticipatory pupil size: Fig. 1I, *F* (1.607, 11.25) = 9.149, *p* = 0.0060, repeated measures one-way ANOVA; Saline v.s. LY341495 10.0 mg/kg: *p* = 0.0240; post-hoc Tukey test; consummatory pupil size: Fig. 1K, *F* (1.612, 11.28) = 9.229 *p* = 0.0058, repeated measures one-way ANOVA; Saline v.s. LY341495 3.0 mg/kg: p =0.0398; Saline v.s. LY341495 10.0 mg/kg: *p* = 0.0254). Effects of LY341495 on pupil size increased along with trial progress (Fig. 1N). Normalized pupil size revealed a dose-dependent attenuation of reward-proximity–related pupillary modulation following LY341495 administration (Fig. 1O). While saline-treated animals exhibited a clear increase in pupil size as reward delivery approached, this modulation was dose-dependently reduced by LY341495, indicating an alteration in the temporal dynamics of pupillary responses associated with reward delivery.

To examine the effects of intraperitoneal injections of LY341495 on gaze direction, we analyzed the central position of the pupil (Fig. 1P,Q). To enable comparisons across mice, the coordinates of the pupil center were calculated relative to the center of the eyelids. Injections of LY341495 did not affect the eye position during the head-fixed temporal conditioning task (X-coordinate: Fig. 1R, *χ²* (2) = 5.250, *p* = 0.079, Friedman test; Fig. 1S, Y-coordinate: *χ²* (2) = 4.750, *p* = 0.120, Friedman test).

### Experiment 2: open-field task in freely-moving mice

To examine the effect of intraperitoneal injections of LY341495 on whole-body motor function irrespective of licking movement, we examined the spontaneous locomotor activity of mice in the open-field box (Fig. 2A-D). Intraperitoneal injections of LY341495 did not affect the spontaneous locomotor activities of the mice (Fig. 2E, *F* (2.141, 17.13) = 1.260, *p* = 0.3109, repeated measures one-way ANOVA). Analysis of the locomotor activity in the time course during the open-field task confirmed that there was no effect of intraperitoneal injections of LY341495 on locomotor activity (Fig. 5F). Intraperitoneal injections of LY341495 have no effect on the time spent in the center area of the open-field box (Fig. 2G,H, *F* (2.220, 17.76) = 1.693, *p* = 0.2110, repeated measures one-way ANOVA).

**Fig. 2:**
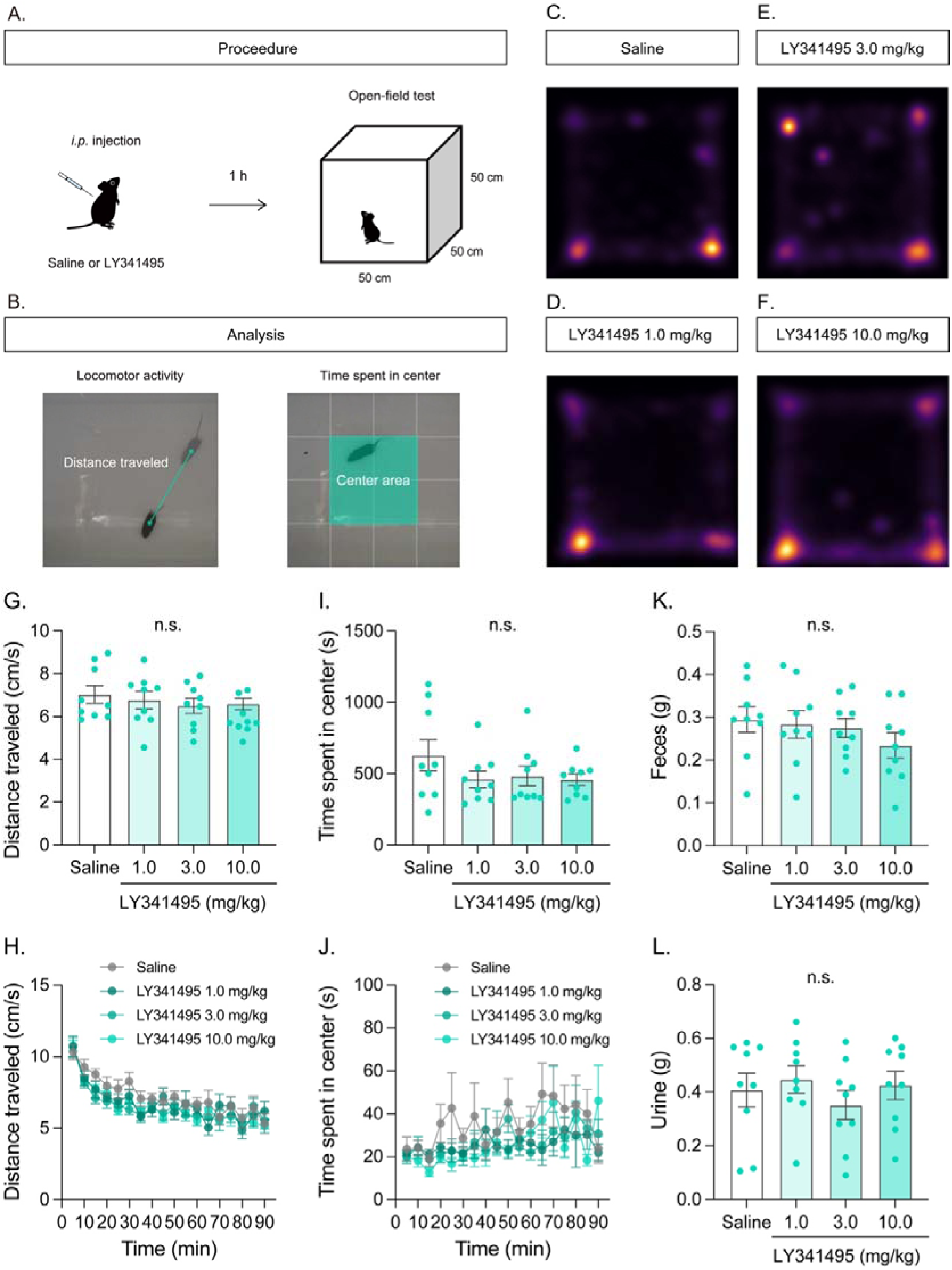
Effects of the intraperitoneal injections of mGluR2/3 receptor antagonist on the performance in the open-field task. A. Experimental timeline of the open-field test, conducted for 90 min, 1 h after intraperitoneal injections of saline or LY341495 (1.0, 3.0, or 10.0 mg/kg). B. Definition of the center area. C-F. Heatmap of the open-field performance in saline (C.), LY341495 1.0 (D.), 3.0 (E.), and 10.0 mg/kg (F.) conditions. G. Overall locomotor activity after the start of the open-field task following saline or LY341495 injection. H. Time course of open-field activity. I. Time spent in the central area of the open-field box after the start of the task. J. Time course of time spent in the central area. The central area is defined as the inner 25 cm × 25 cm portion of the box. K. Fecal output collected after 90 minutes of the open-field task. L. Urine output collected after 90 minutes of the task. Error bars represent standard error of the mean. n.s., not significant. N = 9.

**Fig. 3:**
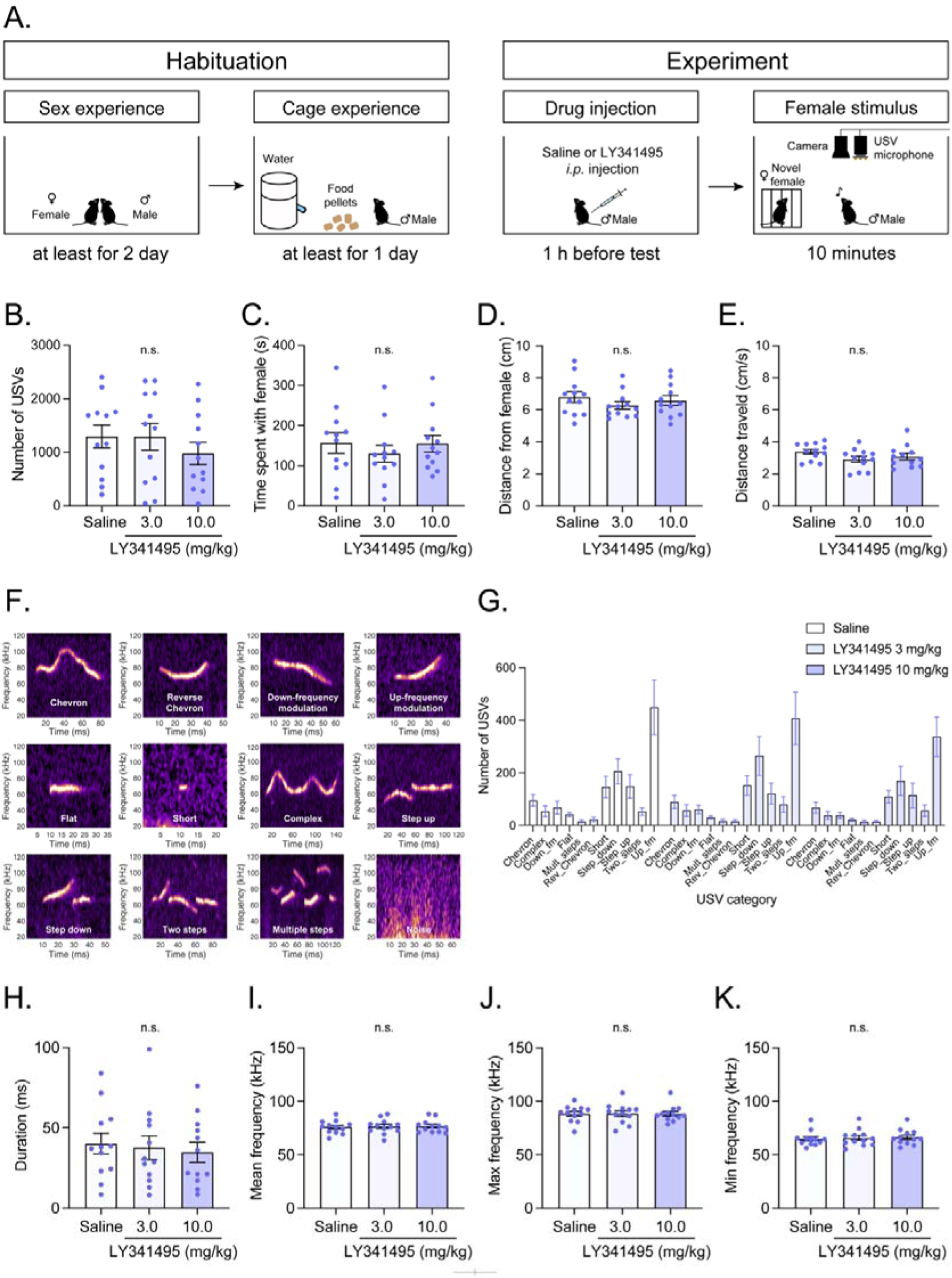
Effects of the intraperitoneal injections of mGluR2/3 receptor antagonist on performance in courtship USVs in male mice. A. Experimental timeline of the USV experiment, conducted 1 h after intraperitoneal injections of saline or LY341495 (3.0, or 10.0 mg/kg). B. Number of USVs. C. Time spent with the female stimulus. D. Distance from female mouse. E. Distance traveled. F. USV classifications G. Number of USVs in each category. Error bars represent standard error of the mean. H. Duration of USV. I. Mean frequency of USV. J . Max frequency of USV. K. Min frequency of USV. \*\**p* < 0.01, n.s., not significant. N = 12.

**Fig. 4:**
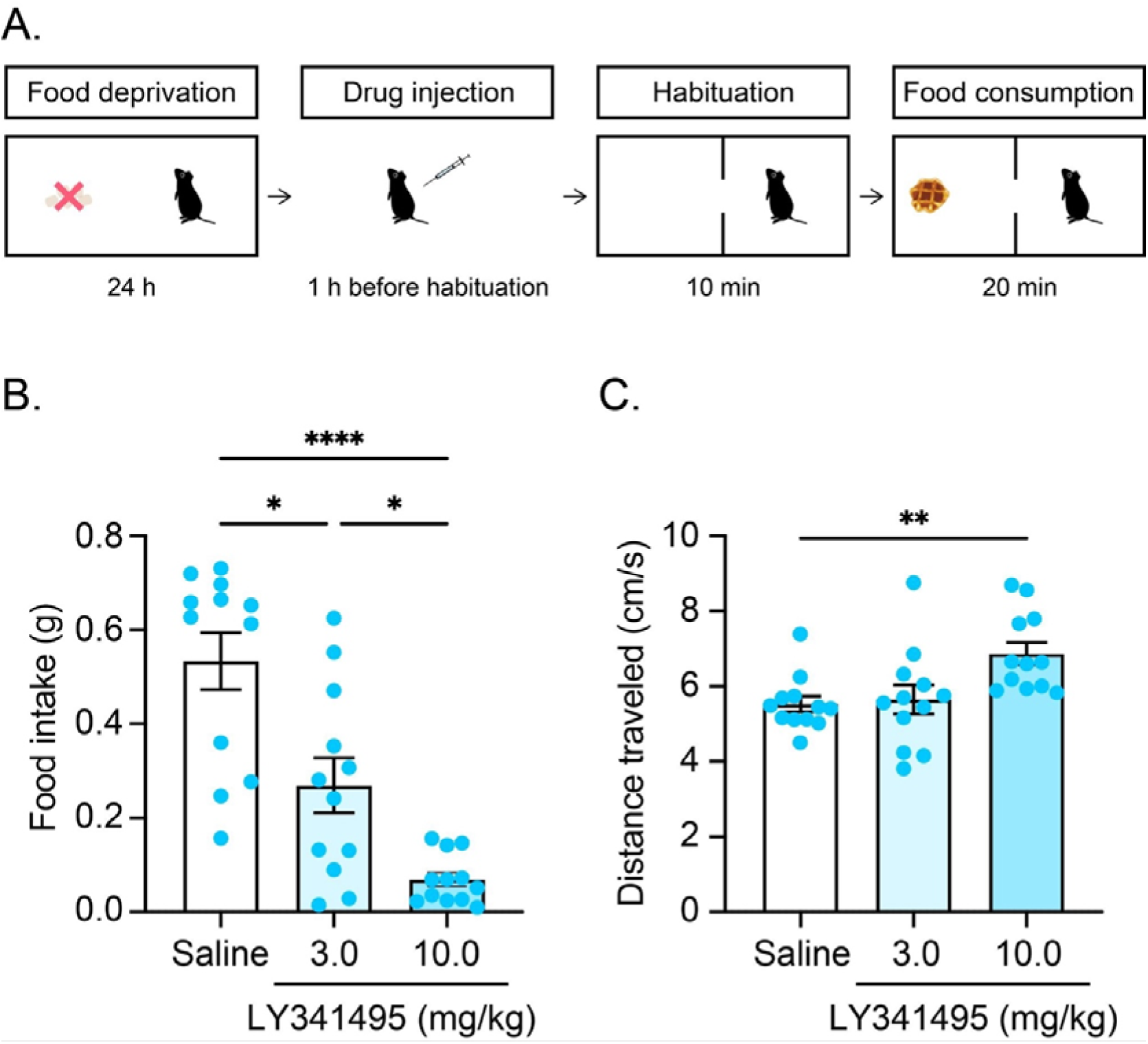
Effects of the intraperitoneal injections of mGluR2/3 receptor antagonist on the performance in the food consumption task. A. Experimental timeline of the food consumption test, conducted for 20 min, 1 h after intraperitoneal injections of saline or LY341495 (1.0, 3.0, or 10.0 mg/kg). B. Food intake during the food consumption task. C. Distance traveled during the food consumption task. Error bars represent standard error of the mean. \**p* < 0.05, \*\**p* < 0.01, \*\*\*\**p* < 0.001, N = 12.

**Fig. 5:**
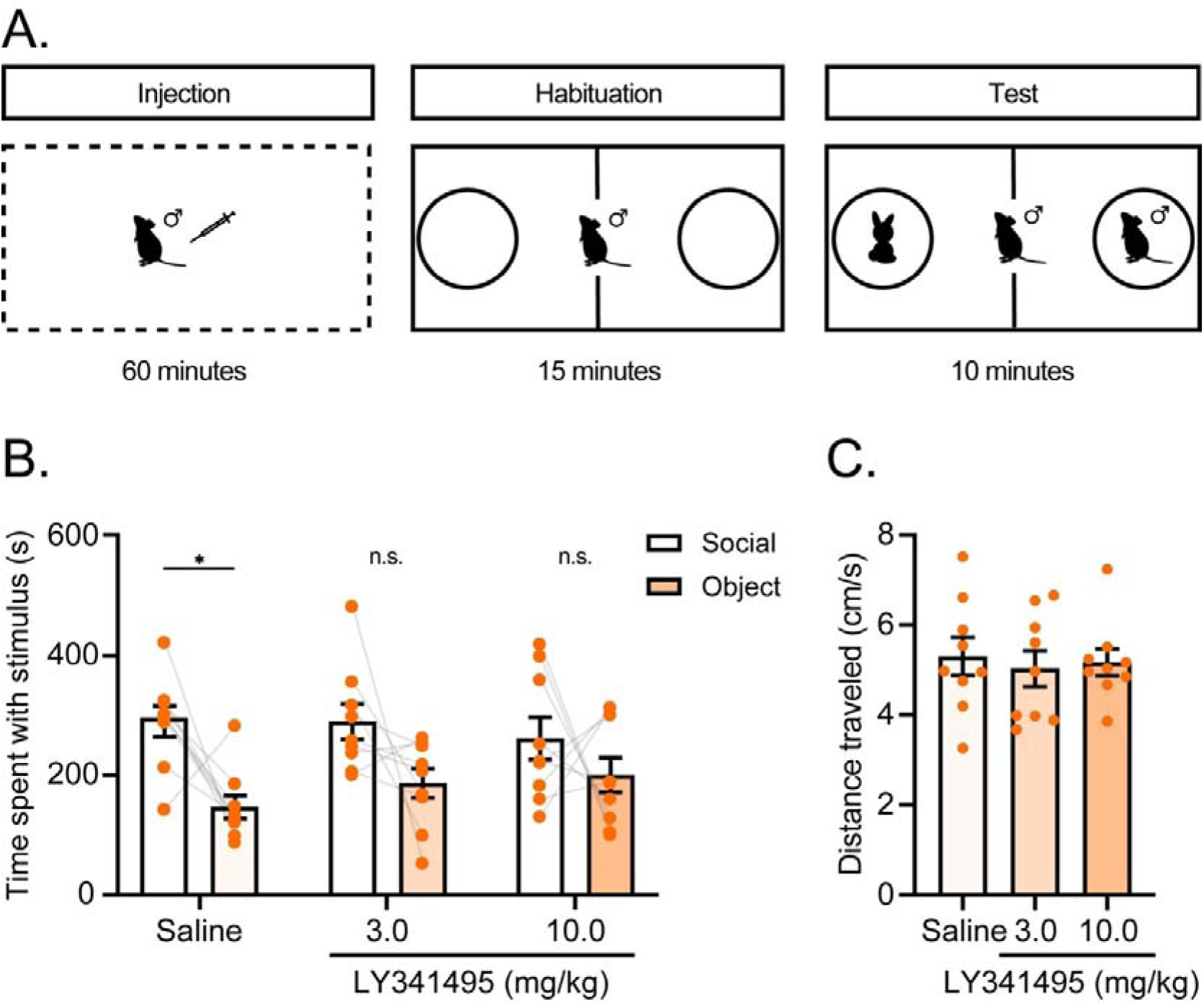
Effects of the intraperitoneal injections of mGluR2/3 receptor antagonist on the performance in the social preference task. A. Experimental procedure of the social preference task. B. Time spent with social and object stimuli. C. Distance traveled during the social preference test. Error bars represent standard error of the mean. \**p* < 0.05, n.s., not significant. N = 9.

To assess the impact of LY341495 on autonomic nervous system function, we collected and quantified measured feces and urine after the mice performed the open-field task. There was no effect of intraperitoneal injection of LY341495 on the amount of feces (Fig. 2I, *F* (2.616, 20.93) = 0.7326, *p* = 0.5269, repeated measures one-way ANOVA) and urine (Fig. 2J, *F* (2.579, 20.64) = 0.7668, *p* = 0.5078, repeated measures one-way ANOVA) collected after the experiment.

### Experiment 3: Courtship interaction task with recording USVs

To assess the effects of LY341495 on orofacial movements, we quantified USVs in male mice exposed to female mice (Fig. 3A). Intraperitoneal administration of LY341495 did not affect courtship vocalizations or behavior, as neither the number of USVs (Fig. 3B, *F* (1.797, 19.77) = 2.269, *p* = 0.1337, repeated measures one-way ANOVA) nor the time spent with females was altered (Fig. 3C, *F* (1.563, 17.19) = 0.7330, *p* = 0.4629, repeated measures one-way ANOVA). Distance from female and locomotor activity were also unaffected (Fig3D, *F* (1.913, 21.04) = 3.155, *p* = 0.0653, repeated measures one-way ANOVA, Fig3E, *F* (1.752, 19.27) = 1.088, *p* = 0.3488, repeated measures one-way ANOVA). In addition, analysis of USV call subtype composition revealed no significant differences between groups (Fig. 3G, main effect of drug: *F* (2,363) = 1.931, *p* = 0.1465; drug × classification interaction: *F* (20,363) = 0.2663, *p* = 0.9995; two-way ANOVA). USVs’ acoustic features were also unaffected (duration: Fig3H, *F* (1.426, 15.69) = 2.407, *p* = 0.1335, repeated measures one-way ANOVA, mean frequency: Fig3I, *F* (1.995, 21.94) = 0.4537, *p* = 0.6406, repeated measures one-way ANOVA, max frequency: Fig3J, *F* (1.965, 21.62) = 0.01861, *p* = 0.9805, repeated measures one-way ANOVA, min frequency: Fig3K, *F* (1.879, 20.66) = 1.533, *p* = 0.2395, repeated measures one-way ANOVA).

### Experiment 4: food consumption task in freely moving mice

To examine the role of mGluR2/3 on feeding behavior, we examined intraperitoneal injections of LY341495 on the freely moving food consumption test. Intraperitoneal injections of LY341495 decreased food consumption in a dose-dependent manner (Fig. 4B, *F* (1.674, 18.42) = 23.71, *p* < 0.0001, repeated measures one-way ANOVA; Saline v.s. LY341495 3 mg/kg, *p* =0.0187, Saline v.s. LY341495 10 mg/kg, *p* < 0.0001, LY341495 3 mg/kg v.s. LY341495 10 mg/kg, *p* = 0.0114, post-hoc Tukey). Locomotor activity while the mice performed the food consumption test was increased by the LY341495 10 mg/kg injection (Fig. 4C, *F* (1.643, 18.08) = 7.322, *p* = 0.0067, repeated measures one-way ANOVA; Saline v.s. LY341495 10 mg/kg, *p* = 0.0068). Even though this was not a quantitative analysis, mice injected with LY341495 did not approach the food.

### Experiment 5: Social preference task

To examine whether mGluR2/3 antagonists affect social-reward –non-food reward– seeking behavior, we conducted a social preference task (Fig. 5A). Under saline conditions, mice showed a preference for the social stimulus over the empty stimulus whereas this preference was not observed following LY341495 administration(Fig. 5B; main effect of drug: *F* (2,24) = 0.6140, *p* = 0.5495; main effect of stimulus: *F* (1,24) = 11.57, *p* = 0.0023, drug × stimulus interaction: *F* (2,24) = 0.6140, *p* = 0.5495, two-way ANOVA; Saline: Social v.s. Object, p = 0.0114, LY341495 3 mg/kg: Social v.s. Object, *p* = 0.0593, LY341495 10 mg/kg: Social v.s. Object, *p* = 0.2523, Social: Saline v.s. LY341495 3 mg/kg, *p* > 0.9999, Saline v.s. LY341495 10 mg/kg, *p* = 0.7484, LY341495 3 mg/kg v.s. LY341495 10 mg/kg, *p* = 0.7945, Object: Saline v.s. LY341495 3 mg/kg, *p* = 0.5726, Saline v.s. LY341495 10 mg/kg, *p* = 0.3678, LY341495 3 mg/kg v.s. LY341495 10 mg/kg, *p* = 0.9344, Tukey’s multiple comparisons test).Locomotor activity were also unaffected (Fig5C; *F* (1.940, 15.52) = 0.1630, *p* = 0.8450, repeated measures one-way ANOVA).

## Discussion

In this study, we investigated the effects of LY341495, a selective mGluR2/3 antagonist, on reward-related behaviors using a combination of head-fixed and freely moving behavioral procedures. In the head-fixed temporal conditioning task, mice exhibited increased anticipatory licking and pupil dilation as the timing of sucrose delivery approached, indicating anticipatory behavioral and physiological responses associated with expected reward delivery. Systemic administration of LY341495 dose-dependently reduced both anticipatory and consummatory licking behaviors. Normalization analyses further demonstrated that LY341495 reduced the overall magnitude of licking behavior without altering its temporal prediction. In addition, LY341495 induced overall pupil dilation and attenuated reward-proximity–related pupillary modulation, suggesting altered arousal or reward processing. To determine whether reduced licking reflected non-specific motor impairments, we assessed spontaneous locomotor activity in a freely moving open-field task. LY341495 did not significantly affect locomotion, center exploration, or excretion, indicating preserved general motor function. Furthermore, to examine whether LY341495 influenced orofacial motor control, we measured USVs during c interaction. LY341495 did not significantly alter USV production, suggesting that LY341495 did not produce a broad impairment of complex orofacial motor behaviors. In contrast, LY341495 dose-dependently reduced food intake in a freely moving feeding task. Moreover, social preference testing revealed that LY341495 reduced social interaction, indicating that the effects of mGluR2/3 antagonism extend beyond food-related rewards to non-food reward processing. These findings demonstrate that mGluR2/3 signaling plays a critical role in regulating reward-seeking behaviors, including both ingestive and social domains, independently of general locomotor or orofacial motor function.

Intraperitoneal injection of LY341495 dose-dependently reduced both anticipatory and consummatory licking in the head-fixed temporal conditioning task, suggesting a critical role for mGluR2/3 in reward-related behaviors. Although LY341495 decreased absolute anticipatory licking, normalized anticipatory licking remained largely unchanged across treatment conditions. This observation indicates that the temporal pattern of anticipatory licking relative to reward delivery was largely preserved despite an overall reduction in licking behavior. However, because LY341495 also reduced consummatory licking and food consumption, the present data do not allow us to determine whether reward prediction itself was unaffected. Therefore, our findings should be interpreted cautiously as suggesting that LY341495 reduced the overall expression of reward-related behavior while leaving its temporal organization relatively intact. Future studies employing neural recordings or behavioral measures that more directly assess reward prediction will be required to determine whether mGluR2/3 antagonism affects predictive processing per se.

LY341495 administration also dose-dependently suppressed feeding behavior in a freely moving food consumption test. This result suggests that the observed disruption of feeding behavior is not attributable to stress induced by the head-fixed experimental context. One possible explanation for this result is that mGluR2/3 inactivation inhibited orofacial movements by affecting motor control functions. However, this motor control explanation appears unlikely. Notably, LY341495 did not reduce spontaneous locomotor activity in the open-field test; instead, locomotion was significantly increased following LY341495 administration. This increase in locomotor activity suggests that general motor function and muscle strength were preserved. In addition, LY341495 did not significantly alter time spent in, or distance traveled through, the center area of the open-field arena. Because center exploration is commonly used as an index of anxiety-like behavior in rodents, these findings suggest that LY341495 did not produce marked anxiogenic or anxiolytic effects under the present experimental conditions. Therefore, altered anxiety levels are unlikely to fully account for the reduction in feeding and licking behavior, supporting the interpretation that LY341495 exerts a relatively specific effect on reward-related or motivational processes. Moreover, the apparent increase in locomotion may partly reflect reduced feeding behavior, as animals typically remain relatively immobile during food consumption. Thus, decreased feeding time could result in a relative increase in locomotor activity.

Another plausible explanation is that mGluR2/3 inactivation altered the motivational value of the sucrose solution. One possibility is that LY341495 reduced the incentive value or rewarding properties of sucrose, thereby decreasing both anticipatory and consummatory licking. Alternatively, the reduction in sucrose consumption could reflect changes in physiological motivational states, such as enhanced satiation or reduced thirst, rather than a direct reduction in reward value. Because the present study did not independently manipulate or measure motivational state, reward valuation, or homeostatic signals, we cannot distinguish between these possibilities. Future studies employing outcome devaluation procedures, progressive ratio schedules, or direct assessments of hunger and thirst states will be necessary to determine whether LY341495 primarily affects reward valuation, motivational drive, or satiety-related processes. Nevertheless, the growing body of evidence that manipulations of mGluR2/3 influence dopaminergic modulation (Swanson et al., 2004; Witkin et al., 2016) and addiction-related behaviors (Moussawi and Kalivas, 2010) supports the broader interpretation that mGluR2/3 signaling contributes to the regulation of reward-related and motivational processes.

Furthermore, LY341495 did not significantly alter courtship USVs during male–female social interaction, suggesting that LY341495 did not produce a broad impairment of orofacial motor control. Although USV production and licking rely on partially distinct motor patterns, the preservation of this complex orofacial behavior argues against a generalized motor deficit as the primary explanation for the reduction in licking behavior. USV production in mice requires coordinated movements of the mouth, larynx, and respiratory systems, and is therefore considered a sensitive index of orofacial motor control. If LY341495 produced a generalized impairment of mouth-related motor output, alterations in USV production would also be expected. However, the absence of significant changes in USVs indicates that the motor commands required for complex orofacial behaviors remained largely intact. This finding further supports the interpretation that the behavioral effects of LY341495 cannot be readily explained by a generalized motor impairment and may instead reflect alterations in reward-related or motivational processes. In addition, these results are consistent with the notion that ingestive behaviors such as licking are influenced not only by motor capacity but also by motivational and reward-related processes that modulate behavioral output.

In contrast, LY341495 reduced social preference in the social interaction task, indicating that mGluR2/3 antagonism affects motivational and reward-related processes beyond food-related reinforcement. Social interaction is widely considered a form of natural reward in rodents, and reductions in social preference are often interpreted as reflecting diminished reward sensitivity or motivational drive. The suppression of social interaction by LY341495 therefore suggests that mGluR2/3 signaling contributes to the regulation of multiple reward domains, including both ingestive and social rewards. This interpretation is consistent with previous studies demonstrating that mGluR2/3 modulation influences dopaminergic transmission and reward-related circuitry, particularly within the mesolimbic pathway (Swanson et al., 2004; Witkin et al., 2016). Importantly, because locomotor activity and orofacial motor function were preserved, the reduction in social preference is unlikely to be explained by general motor impairment. Instead, these findings support the idea that mGluR2/3 signaling plays a broader role in regulating motivational processes underlying reward-seeking behavior across distinct behavioral domains. Importantly, the observation that LY341495 affected both ingestive and social reward-related behaviors may help constrain potential neural mechanisms underlying these effects. Although the present study did not identify the specific brain regions involved, the convergence of effects across distinct reward modalities suggests that mGluR2/3 signaling may influence neural systems that contribute broadly to motivational and reward-related processing rather than circuits exclusively dedicated to feeding behavior. This interpretation is consistent with the widespread expression of mGluR2/3 receptors throughout reward-related networks, including the prefrontal cortex, nucleus accumbens, amygdala, and hypothalamus. Nevertheless, because the present study relied on systemic drug administration, the specific neural circuits mediating these behavioral effects remain unresolved and will require future investigation using region-specific and circuit-specific approaches.

Intraperitoneal injection of LY341495 induced significant, dose-dependent pupil dilation, implying increased sympathetic nervous system activity or decreased parasympathetic nervous system activity. Although the precise mechanism remains unclear, elevated extracellular dopamine levels following LY341495 administration (Witkin et al., 2016) may contribute to the observed pupillary response. Because pupil diameter is commonly used as an index of arousal state, this finding may indicate that mGluR2/3 blockade alters central arousal mechanisms in addition to reward-related processes. Although LY341495 did not affect fecal output or urination, the observed pupil dilation suggests a possible influence of mGluR2/3 blockade on autonomic function. Because different autonomic measures may reflect distinct physiological processes, the apparent dissociation between pupillary and excretory responses suggests that LY341495 may selectively affect specific autonomic pathways. A microdialysis study demonstrated increased dopamine release in the nucleus accumbens and prefrontal cortex approximately 60 minutes after LY341495 administration (Witkin et al., 2016), a time course similar to that used in the present study. Because dopaminergic signaling has been implicated in pupil dilation and arousal regulation, elevated dopamine transmission may contribute to the observed pupillary response. However, because LY341495 was administered systemically, the present study cannot determine whether the observed behavioral and autonomic effects were mediated primarily by central or peripheral mechanisms. In particular, the pupil dilation observed following LY341495 administration may reflect direct or indirect influences on peripheral autonomic pathways in addition to central neural circuits. Future studies combining central administration approaches, neural recordings, and physiological monitoring will be necessary to dissociate central and peripheral contributions to the observed effects. Importantly, despite these physiological changes, LY341495 did not impair spontaneous locomotor activity and did not alter USV production. These findings support the interpretation that mGluR2/3 antagonism primarily affects motivational and reward-related processes rather than producing broad motor impairment.

The present findings may have broader implications for understanding how mGluR2/3 signaling contributes to reward-related behavior. Although manipulations of dopaminergic systems often influence both reward processing and locomotor activity, LY341495 reduced ingestive and social reward-related behaviors without producing clear motor deficits. This dissociation suggests that mGluR2/3 signaling may contribute to motivational and reward-related processes that are at least partially separable from general motor function. Future studies combining behavioral, physiological, and neural recording approaches will be necessary to determine how mGluR2/3 signaling interacts with dopaminergic and other neuromodulatory systems to regulate reward-seeking behavior.

Our study has several limitations. First, because LY341495 was administered systemically, we cannot determine whether the observed behavioral and physiological effects were mediated primarily by central or peripheral mechanisms. This limitation is particularly important for the interpretation of pupil dilation and reward-related behavioral changes, as both central neural circuits and peripheral autonomic pathways may contribute to these effects. Therefore, although our findings are consistent with a role for mGluR2/3 signaling in reward-related processes, direct evidence linking the observed behavioral effects to specific central neural mechanisms remains lacking. Future research should explore the effects of intracerebroventricular injections of mGluR2/3 antagonists, combined with heart rate and respiration monitoring, to differentiate their impacts on the central and peripheral nervous systems. Second, although we assessed locomotor activity and USVs to evaluate potential motor impairments, we did not perform a comprehensive analysis of fine-scale motor patterns, including detailed orofacial movements, posture, or micro-motor behaviors. While the preservation of USV production argues against a broad orofacial motor deficit, it remains possible that subtle motor changes not captured by our measurements contributed to the observed behavioral effects. Future studies using high-resolution video analysis and machine learning–based behavioral classification could provide a more detailed characterization of motor and behavioral changes following mGluR2/3 manipulation. Third, although LY341495 reduced social preference, we did not directly distinguish whether this effect reflected reduced social motivation, altered anxiety levels, or changes in social cognition. Because social interaction behavior is influenced by multiple psychological processes, including anxiety, novelty processing, and exploratory drive, additional behavioral paradigms will be necessary to clarify the specific mechanisms underlying the observed reduction in social interaction. Fourth, we did not identify the specific brain regions or neural pathways affected by systemic LY341495 administration. Given that mGluR2/3 receptors are widely distributed across reward-related brain regions, including the prefrontal cortex, nucleus accumbens, amygdala, and hypothalamus, region-specific manipulations using local pharmacological injections or circuit-specific approaches will be crucial for elucidating the neural mechanisms underlying the regulation of ingestive and social reward behaviors. Fifth, only male mice were included in the present study. Therefore, it remains unclear whether the behavioral and physiological effects of LY341495 observed here generalize to females. Given the well-documented sex differences in reward processing, social behavior, and glutamatergic signaling, future studies should examine the effects of mGluR2/3 antagonism in female animals and directly compare responses between sexes. Addressing these limitations in future studies will help clarify the neural and psychological mechanisms by which mGluR2/3 signaling regulates reward-related behaviors.

In conclusion, our findings demonstrate that systemic administration of the mGluR2/3 antagonist LY341495 dose-dependently disrupted reward-related behaviors, including anticipatory and consummatory licking, food intake, and social preference, while leaving general locomotion and USVs largely intact. These results suggest that mGluR2/3 signaling plays a critical role in regulating reward-related motivation and consumption rather than broadly impairing motor function. Although mGluR2/3 receptors have previously been implicated in learning, memory, and affective processes, their causal involvement in reward-related behavior—particularly ingestive behavior—has remained unclear. Our findings indicate that mGluR2/3 contributes to the modulation of reward-driven behavior across both food-related and social domains. Furthermore, the preservation of the temporal organization of anticipatory licking despite an overall reduction in behavioral output suggests that mGluR2/3 signaling may influence the expression of reward-related behavior rather than producing a complete disruption of temporal reward-related responding. These results provide new insights into the psychological and neurobiological mechanisms underlying reward-seeking and consumption behaviors. Given the broad role of reward processing in conditions such as obesity, eating disorders, and motivational deficits, mGluR2/3 signaling may represent a promising target for understanding and potentially treating disorders characterized by dysregulated reward-related behaviors.

## Declarations

### Ethics approval and consent to participate

The experimental and housing protocols adhered to the Japanese National Regulations for Animal Welfare and were approved by the Animal Care and Use Committee of the Keio University.

## Availability of data and materials

Data supporting the findings of this study are available from the corresponding author upon reasonable request. The original codes written for the analysis are available from the corresponding author upon reasonable request.

## Authors’ contributions

HI and KT designed the experiments. HI collected the data of the head-fixed temporal conditioning experiment with the help of MY, SM, KH, and KT. HI, MY, and KT collected the data of the open-field test. HI and MY collected the data of the social interaction task with recording USVs with the help of YT. HI, MY, YT, and KT analyzed the data with the help of KY. HI, MY, and KT wrote the manuscript. HI, MY, YT, and KT created all figures. HI, MY, YT, SM, KY, KH, and KT discussed the data and commented on the manuscript. KT revised the manuscript accordingly.

## Acknowledgement

We thank Haruki Kasahara, Shunsuke Nakajima, Shudo Yoshida, and Rei Takahashi for their assistance on animal care. The authors used ChatGPT (OpenAI) to improve the clarity and readability of the manuscript. All authors reviewed and approved the final version of the manuscript and take full responsibility for its content.

